# Plasma membrane-endoplasmic reticulum coupling probed with genetically-encoded voltage sensors

**DOI:** 10.1101/2025.10.03.680358

**Authors:** Masoud Sepehri Rad, Meyer B. Jackson

**Affiliations:** Department of Neuroscience, University of Wisconsin –Madison, Madison WI 53705

## Abstract

The endoplasmic reticulum (ER) forms an elaborate contiguous network extending through the cytoplasm of eukaryotic cells. The ER is surrounded by a membrane that separates its lumen from the cytoplasm. The ER membrane harbors channels and pumps capable of controlling ion flux and creating a voltage gradient. Because the ER membrane potential is difficult to study experimentally little is known about how voltage influences its many vital functions. Here we introduce optical probes of ER membrane potential derived from the hybrid voltage sensor (hVoS) family of genetically-encoded voltage sensors. Probes were targeted to the ER using motifs from three ER proteins, Sec61β, cytochrome P450, and cytochrome b_5_ type A. As shown recently with other types of ER voltage sensors, patch-clamp fluorometry recording with our new probes demonstrated that voltage steps applied to the plasma membrane elicit a voltage change at the ER membrane. These probes exhibited subtle differences in their responses suggesting they target different ER compartments. The steeper voltage dependence of Sec61β-hVoS (mCerulean3-Sec61β) signals suggested that this probe targets an ER compartment rich in voltage-gated ion channels. The ER voltage change is slow, but its onset is virtually synchronous with the plasma membrane voltage step. This suggests a direct electrical coupling into the ER lumen through plasma membrane-ER contacts. Analysis with the aid of an equivalent circuit provided an estimate of the resistance of these contacts. The rapid, direct transmission of voltage changes from the plasma membrane to the ER provides a mechanism for regulating ER function that could be especially important in excitable cells. The sensors introduced here provide researchers with powerful tools for imaging ER voltage and assessing its impact on cellular function.

**Significance:** The ER is encompassed by a membrane, and the voltage across this membrane is likely to play important roles in many cellular functions. It is difficult to study this voltage due to the inaccessibility of organelle membranes. We introduced new hybrid genetically-encoded optical voltage sensors that target the ER. These probes showed that a voltage step applied to the plasma membrane changes the voltage at the ER membrane in a manner consistent with direct electrical coupling to the cell surface through contacts. This coupling will enable voltage changes at the plasma membrane to influence ER function. The probes introduced here will enable researchers to image ER voltage and probe plasma membrane-ER signaling.

## Introduction

The ER carries out many vital functions including protein translation, processing, and sorting, as well as lipid biosynthesis and cytosolic Ca^2+^ regulation (1, 2). ER dysfunction is associated with a variety of pathological conditions, including lipid storage diseases, neuropathy, neurodegeneration, and cancer (3, 4). Dynamic interactions with the plasma membrane, Golgi, mitochondria, nucleus, and other structures enable the ER to serve as a critical hub in cellular control (5, 6). The ER generates rises and falls in cytosolic Ca^2+^ through release and sequestration (7), and ER Ca^2+^ signaling in neurons regulates synaptic transmission and plasticity (8-10). The complex biphasic nature of Ca^2+^ release enables the ER to generate Ca^2+^ oscillations (11), and their frequency serves as a signal in regulating gene expression (12). The ER is formed by a membrane that harbors many ion channels (13). These channels presumably establish a voltage gradient across the ER membrane, Ψ_ER_, that will drive the flux of charged permeable species. Because it is experimentally inaccessible little is known about how Ψ_ER_ is controlled, and how it impacts cellular function.

Organelle membranes cannot be studied with typical electrodes used in conventional electrophysiology, but a genetically encoded voltage indicator (GEVI) could overcome this problem by targeting a specific organelle for optical recording of its voltage (14). ER targeted GEVIs have been used to provide estimates of resting Ψ_ER_ (14, 15). A probe derived by mutating the voltage sensing domain of the GEVI ArcLight produced signals originating partially from the ER (16). This study indicated that voltage steps applied to the plasma membrane produced changes in Ψ_ER_. Localized unquenching of a rhodamine-based synthetic voltage sensor in the ER confirmed the coupling of the plasma membrane to the ER and further showed that depolarizing the plasma membrane made the cytoplasm more positive with respect to the ER lumen (17). A probe generated by incorporating an ER-retention motif into the GEVI ASAP3 also recorded changes in Ψ_ER_ and characterized responses to a variety of signals known to act on the ER (18). These various probes illustrate the enormous potential of ER-GEVIs to explore how Ψ_ER_ responds to biological signals and how it contributes to ER functions.

Probes of Ψ_ER_ have all revealed a surprisingly rapid coupling between the plasma membrane and ER. Voltage steps to the plasma membrane elicit clear changes in Ψ_ER_ within roughly 100-200 msec (16-18). In the present study we introduced a new family of ER-GEVIs and used these probes to investigate this coupling. Our ER-GEVIs were derived from hybrid voltage sensors (hVoS) (19), which have been widely employed in the study of voltage changes at the plasma membrane associated with neuronal excitation (20-22). hVoS probes consist of a fluorescent protein tagged with a membrane targeting motif. Their voltage sensitivity depends on the presence of a second molecule, dipicrylamine (DPA), which partitions into lipid bilayers. Voltage drives negatively charged DPA between the two faces of a lipid bilayer, and a Förster resonance energy transfer (FRET) interaction between DPA and the fluorescent protein converts this movement into a voltage dependent optical signal. The general ease of targeting fluorescent proteins to organelles (23) greatly facilitates the subcellular targeting of hVoS probes, as was demonstrated by the development of a lysosomal voltage sensor (15). Here we report hVoS-based ER-GEVIs that target the ER using motifs from three ER proteins. Sec61β is found primarily in the rough ER where it functions in precursor translocation (24, 25). Cytochrome P450 (P450) is restricted to the ER (26) by its N-terminus (27). Cytochrome b_5_ type A (Cb5) targets the ER through its C-terminus (28-30). When expressed in HEK293T cells each of these probes targeted the ER with high specificity, and were effectively excluded from the plasma membrane. In patch-clamped HEK293T cells these probes generated robust fluorescence changes in response to voltage steps applied to the plasma membrane. Our experiments confirm the rapid coupling and positive changes in Ψ_ER_ upon plasma membrane depolarization. We analyzed the kinetics of the optical signal with the aid of an equivalent circuit to characterize the conductive nature of plasma membrane-ER contacts. This work establishes direct electrical coupling with the plasma membrane as a signaling mechanism that modulates Ψ_ER_. The powerful optical tools introduced here can be used to investigate Ψ_ER_ in diverse ER compartments in any cell of interest, and to monitor how Ψ_ER_ responds to biological signals.

## Materials and Methods

### Plasmid and DNA constructs

The plasma membrane targeted hVoS probe was generated as described previously (19, 31) and is a slight modification of Addgene plasmid #45281 where the fluorescent protein was upgraded to mCerulean3 fluorescent protein (mCeFP) (32) to generate hVoS3. The construct Bongwoori-Pos6 was a gift from Bradley Baker (Addgene plasmid # 101655) (33). mCherry-Sec61β was Addgene plasmid # 49155 (34) and Lck-mScarlet was Addgene plasmid #98821 (23).

We constructed P450-mCeFP as follows. Step 1: Primers for amplification of the fluorescent protein in the first PCR reaction were: 5-CTCTGGAAACAGAGCTATGGGGGAGGgatggtgagcaagggcgag-3 and CE-49: 5-ACGGGCCCTCTAGACTCGAGTTCTAGATCAgccgagagtgatcccggc-3. Step 2: Using the step 1 product as a template, we then used primers: 5-GTGCTGGGGCTCTGTCTCTCCTGTTTGCTTCTCCTTTCACTCTGGAAACAGAGCTATGGG -3 and CE-49 to append the mCeFP to the C-terminus of the ER targeting motif (35). Step 3: We used the step 2 product as a template with primers: 5-GGGAGACCCAAGCTGGCTAGCACCATGGACCCTGTGGTGGTGCTGGGGCTCTGTCTCTCC-3 and CE-49 to introduce Nhe1 and Xho1 sites at the N-terminus of P450 and C-terminus of mCeFP respectively. The step 3 PCR product was then digested with Nhe1 and Xho1 and inserted into the corresponding sites of the Bongwoori-Pos6 construct to generate P450-mCeFP.

We constructed Cb5-mCeFP as follows. Step 1: The primers used for amplification of the fluorescent protein were: 5-GCCCTGGTGGTGGCCCTGATGTACCGCCTGTACATGGCCGAGGACatggtgagcaagggc-3 and CE-49: 5-ACGGGCCCTCTAGACTCGAGTTCTAGATCAgccgagagtgatcccggc-3. Step 2: The first PCR product was used as a template with the primers: 5-TCCAACTCCTCCTGGTGGACCAACTGGGTGATCCCCGCCATCTCCGCCCTGGTGGTGGCC-3 and CE-49. mCeFP was fused to the C-terminus of the ER targeting motif (36). Step 3: The step 2 product was used as a template with primers: 5-ATAGGGAGACCCAAGCTGGCTAGCACCATGATCACCACCGTGGAGTCCAACTCCTCCTGG-3 and CE-49 to introduce Nhe1 and Xho1 sites at the N-terminus of the ER targeting motif and the C-terminus of mCeFP respectively. Step 4: The step 3 product was then digested with Nhe1 and Xho1 and inserted into the corresponding sites of the Bongwoori-pos6 construct to generate the Cb5-mCeFP construct. Then the mCeFP-Cb5 plasmid was generated as follows. Step 1: The primers used for amplification of the fluorescent protein were: CE-67: 5-GGGAGACCCAAGCTGGCTAGCACCatggtgagcaagggcgaggagctg-3 and: 5-GGACTCCACGGTGGTGATgccgagagtgatcccggc-3. We used Cb5-mCeFP as a template to amplify the ER targeting motif with primers: 5-gccgggatcactctcggcATCACCACCGTGGAGTCC-3 and CB-109: 5-ACGGGCCCTCTAGACTCGAGTTCTAGATCAGTCCTCGGCCATGTACAG-3. In the second PCR step, we used primers CE-67 and CB-109 and combined the first step PCR products. The second step PCR product then was digested with Nhe1 and Xho1 and inserted into the corresponding sites of the Bongwoori-Pos6 construct

Finally, we constructed the mCeFP-Sec61β plasmid as follows. Step 1: The primers used for amplification of the fluorescent protein were: CE-67: 5-GGGAGACCCAAGCTGGCTAGCACCatggtgagcaagggcgaggagctg-3 and: 5-cggaccaggcatagatctgccgagagtgatcccggc-3. We used mCherry-Sec61β as a template to amplify the ER targeting motif with primers: 5-gccgggatcactctcggcagatctatgcctggtccg-3 and SE-106: 5-ACGGGCCCTCTAGACTCGAGTTCTAGATCAcgaacgagtgtacttgcc-3. In the second PCR step, we used primers CE-67 and SE-106 and combined the PCR products of the first step. The second step PCR product then was digested with restriction enzymes Nhe1 and Xho1 and inserted into the corresponding sites of the Bongwoori-Pos6 construct.

DNA sequences for all of constructs were confirmed by DNA sequence analysis using the dye-termination method (Functional Biosciences, Inc).

### Cell culture

HEK293T cells were cultured in high glucose DMEM (Gibco) supplemented with 10% fetal bovine serum (Invitrogen) and plated on cover slips coated with poly-L-lysine (Sigma). Cultures were maintained in 24-well plates and incubated at 37 °C in 5% CO_2_. Cells were transfected using Lipofectamine 2000 (Invitrogen) according to the manufacturer’s instructions.

### Patch clamp recording

Cells were patch clamped with an Axopatch 200B amplifier (Molecular Devices) controlled by PCLAMP software through a Digidata 1440 interface (Molecular Devices). During recordings cells were bathed at room temperature in a solution consisting of 150 mM NaCl, 4 mM KCl, 2 mM CaCl_2_, 1 mM MgCl_2_, 5 mM D-glucose, and 5 mM HEPES, pH 7.4, with the addition of 4 μM DPA unless otherwise noted. Patch pipettes (resistance 3-5 MΩ) were filled with a solution consisting of 120 mM K-aspartate, 4 mM NaCl, 4 mM MgCl_2_, 1 mM CaCl_2_, 10 mM EGTA, 3 mM Na_2_ATP, and 5 mM HEPES, pH 7.2. Cell capacitance and series resistance were determined by cancelation of the transient charging current during a stepwise voltage change (37).

### Voltage imaging

Patch clamped cells were viewed with an Olympus BX51 microscope equipped with a DaVinci 2K CMOS camera (SciMeasure Imaging). An LED with peak emission at 435 nm (Prizmatix) supplied illumination and a CFP filter cube selected bands for excitation and emission. Images were acquired using an Olympus XLUMPlanFl 10X objective (NA = 0.6). Image acquisition was controlled by Turbo-SM software, provided with the camera. All fluorescence traces displayed are averages over regions encompassing an entire cell, except for Fig. 3 where the regions are indicated in the images.

### Probe localization

HEK293T cells expressing fluorescent probes were imaged with a Nikon A1R Ti2 confocal microscope equipped with a Plan-Apo 60x objective (NA 1.40). mCeFP was excited with a 405 nm laser and a 425-475 nm band-pass filter was used for emission. To excite mCherry-Sec61β and lck-mScarlet a 561 nm laser was used along with a 570-620 nm band-pass emission filter. Images were acquired and analyzed with the computer program NIS-Elements Viewer (Nikon), which was used to calculate the Pearson’s correlation coefficient between the signals from the two channels within an area encompassing a selected cell.

### Analysis

GEVI signals were analyzed with Turbo-SM to select regions of interest. Dark frames were recorded in the absence of illumination prior to acquisition of each data episode, and were subtracted pixel-by-pixel from recorded data. Dark readings were generally <0.2% of the fluorescence readings from cells. ΔF/F was calculated as (F-F_0_)/F_0_, with F_0_ taken as the ten frame average from the same region of interest acquired before a voltage step. Data was exported to Origin 2023b (OriginLab Corporation) for further analysis, including baseline subtraction and fitting to Boltzmann and exponential functions.

## Results

### Localization

We evaluated the targeting of ER-GEVI constructs using confocal microscopy. P450-mCeFP targets ER with the N-terminal 27 amino acids derived from cytochrome P450 (27). Cells co-transfected with P450-mCeFP and the plasma membrane label lck-mScarlet (23) express these proteins in distinct locations, with mCeFP fluorescence distributed through the cytoplasm (Fig. 1A) and mScarlet fluorescence restricted to the cell surface (Fig. 1B). The merged image shows clear separation of the two labels (Fig. 1C). A line scan through the cell provides another illustration of the different distributions of the ER and plasma membrane probes, and indicates that P450-mCeFP is effectively excluded from the plasma membrane (Fig. 1D). This is further supported by the low Pearson’s correlation coefficient computed over the entire cell between the two probes of 0.23, (Fig. 1I). Two additional constructs, mCeFP-Sec61β and mCeFP-Cb5, were evaluated in the same way. Both of these probes also targeted the cytoplasm and were excluded from the plasma membrane. Exclusion from the plasma membrane was again assessed with the Pearson’s correlation coefficient with lck-mScarlet and the values were similar to those seen with P450-mCeFP (Fig. 1I). We also examined the colocalization of P450-mCeFP and mCeFP-Cb5 with the widely used ER label mCherry-Sec61β. In a cell expressing P450-CeFP and mCherry-Sec61β, both probes had similar cytoplasmic distributions (Figs. 1E and 1F). The merged image showed very high overlap (Fig. 1G), and the line scans showed similar spatial distributions with good registration between the peaks and troughs (Fig. 1H).

**Figure 1.**
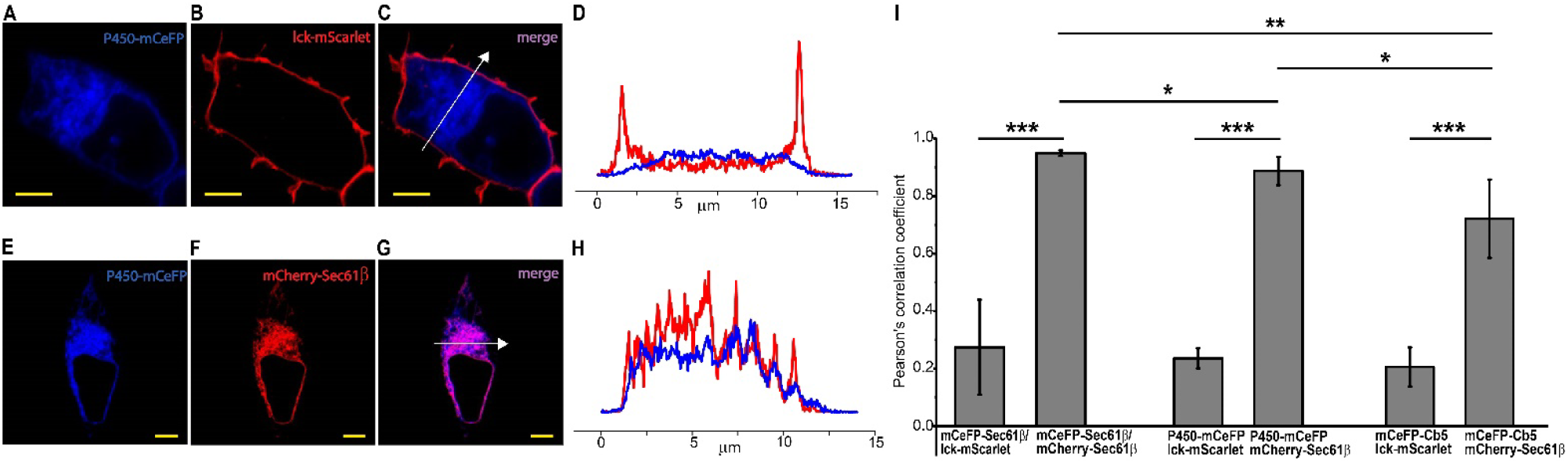
Probe localization. **A.** Confocal fluorescence image of P450-mCeFP in an HEK293T cell. **B**. Fluorescence of the red plasma membrane label (lck-mScarlet) in the same cell as A. **C**. Merging of images A and B reveals non-overlapping distributions between P450-mCeFP and lck-mScarlet. **D**. Scan through the line indicated in C compares the two distributions. **E**. Fluorescence of P450-mCeFP. **F**. Fluorescence of mCherry-Sec61β. **G**. Merging of images E and F reveals the colocalization of both probes as purple. **H**. Scan through the line indicated in G compares the two distributions. **I**. Colocalization of the three ER constructs with mCherry-Sec61β and lck-mScarlet was evaluated within entire cells (5 cells for each pair of labels) with Pearson’s correlation coefficients. Plots show means ± standard errors; P < 0.0001 for all mCeFP-ER::lck-mScarlet versus respective mCeFP-ER::mCherry-Sec61β comparisons (^***^). P = 0.0055 for mCeFP-Sec61β::mCherry-Sec61β versus mCeFP-Cb5:: mCherry-Sec61β (^**^); P = 0.02164 for mCeFP-Sec61β::mCherry-Sec61β versus P450-mCeFP::mCherry-Sec61β (^*^); P = 0.03351 for mP450-CeFP::mCherry-Sec61β versus mCeFP-Cb5::mCherry-Sec61β (^*^). Scale bars = 5 μm in all images.

To compare our ER probes we first note that colocalization of Sec61β with two different tags (mCeFP-Sec61β and mCherry-Sec61β) was very high; the Pearson’s correlation coefficient was 0.95 (Fig. 1I). This assessment between two Sec61β probes essentially evaluates our method for quantifying colocalization. When we turned to the colocalization of mCherry-Sec61β with P450-mCeFP and with mCeFP-Cb5, we found significantly lower correlation coefficients of 0.89 and 0.72, respectively (Fig. 1I). Although this correlation is still high, the comparisons between each of the three pairs suggest that the probes exhibit subtle differences in their distributions, with small amounts of preferential targeting. Either they target different compartments within the ER, or they exhibit different degrees of spillover to another membranous organelle such as the Golgi. Subtle differences in the localization of these probes within the ER are not unexpected given the functions of the parent proteins. P450 and Cb5 reside in both rough and smooth ER, while Sec61β resides only in rough ER where it functions in protein synthesis (25, 26). In summary, all three of our candidate ER-GEVIs were excluded from the plasma membrane very effectively. They target the ER but appear to show subtle differences in their distribution within the ER and possibly between other internal structures. These small intra-ER targeting differences are consistent with subtle differences in the Ψ_ER_ sensing properties of these probes presented below.

### Optical recording of ER voltage changes

To test the ability of our candidate ER-GEVIs to sense voltage we performed patch-clamp fluorometry. In a cell expressing mCeFP-Sec61β (Fig. 2A) probe fluorescence can be seen distributed throughout the cell (Fig. 2B), and voltage steps from -70 to -200 mV elicited large and clear fluorescence increases. Responses to other voltages will be presented below (Fig. 4), but for our initial characterizations we optimized response size by using pulses from -70 to -200 mV. The fluorescence begins to change at the onset of the voltage step and continues over the two second duration of the pulse (Fig. 2C, olive trace, voltage indicated by the black trace below). The fluorescence change depended on the presence of DPA (4 μM): when DPA was omitted the fluorescence remained flat as the voltage changed (Fig. 2C, light green trace), indicating that the signal tracks DPA movement within the membrane. Thus, the signal reports a change in Ψ_ER_ and not some other cellular response to voltage such as a change in pH that might alter mCeFP fluorescence. The slow change in fluorescence to a stepwise change in voltage is a distinct feature of ER-GEVIs. We expressed the plasma membrane probe hVoS3 in an HEK293T cell (Fig. 2D). Resting fluorescence highlights the plasma membrane (Fig. 2E) and a voltage step produced a rapid fluorescence change (Fig. 2F). This is consistent with the established rapid movement of DPA within membranes (38), and the rapid kinetics of plasma membrane hVoS responses settling in < 1 msec (19, 31). Thus, the time course of the response of mCeFP-Sec61β to voltage reflects the charging kinetics of the ER membrane.

**Figure 2.**
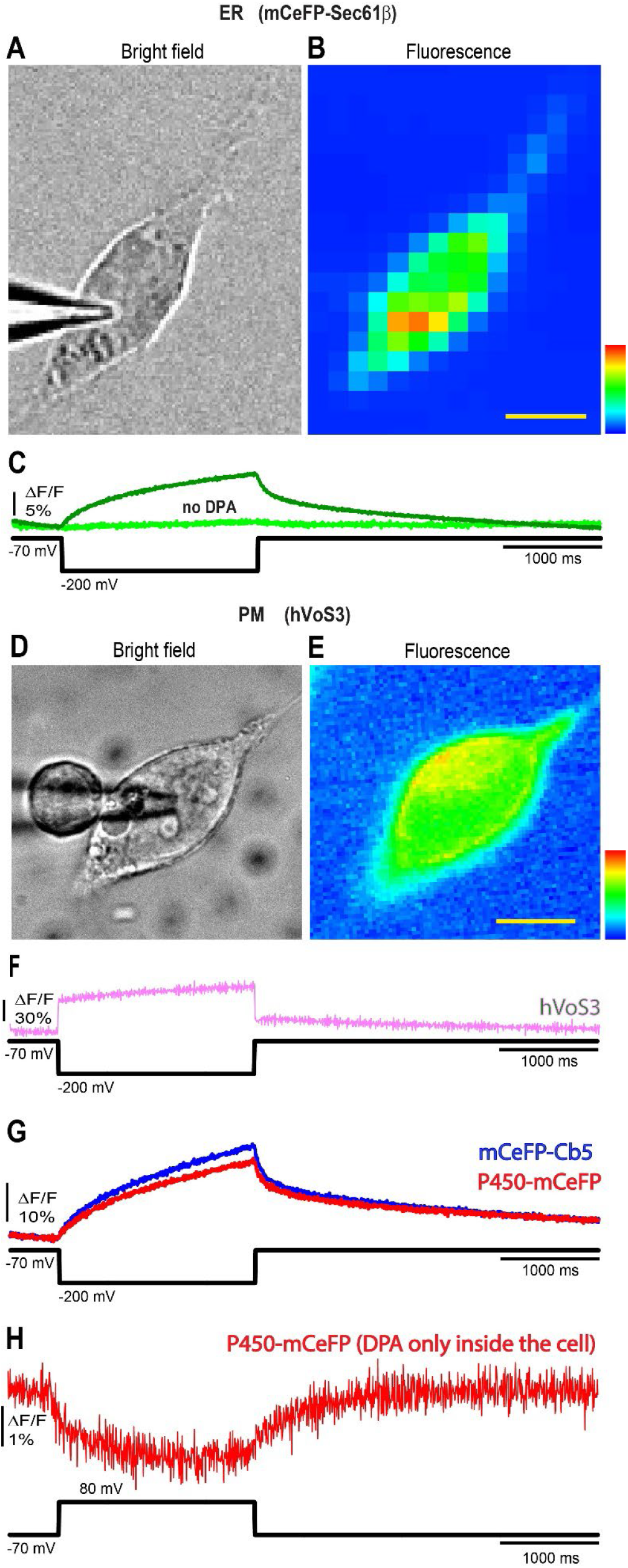
ER-GEVI and hVoS3 signals in voltage-clamped HEK293T cells. **A.** Bright field image of a patch-clamped HEK293T cell expressing mCeFP-Sec61β with patch electrode visible to the left. **B**. Fluorescence image of the same cell. **C**. Traces of fluorescence versus time as voltage steps were applied from -70 mV to -200 mV (indicated by the black trace below) with DPA (dark green trace) and without DPA (light green trace) in the bath solution. **D**. Bright field and **E**. fluorescence image of a cell expressing the plasma membrane probe hVoS3. **F**. Fluorescence versus time illustrates the much faster response to a voltage step from -70 to -200 mV. **G**. Fluorescence versus time as a voltage step was applied from -70 mV to -200 mV (indicated by the black trace below) to a patch-clamped HEK293T cell expressing mCeFP-Cb5 (blue) or P450-mCeFP (red). **H**. Fluorescence versus time in a P450-mCeFP expressing HEK293T cell in a DPA-free bathing solution. The cell was first incubated 30 minutes in 8 μM DPA and then transferred to DPA-free for recording. The recording was made within 10 minutes of transfer. The voltage was stepped from -70 to +80 mV. No temporal filtering was used. In all traces baselines were fitted to exponentials and subtracted. Traces were recorded at a frame rate of 200 Hz. Scale bar = 10 μm.

Probes built with ER targeting motifs derived from two other ER proteins, P450 and Cb5, also produced clear fluorescence changes in response to voltage steps (Fig. 2G). Like the signals from mCeFP-Sec61β, the signals from mCeFP-Cb5 and P450-mCeFP start with the voltage step and continue for the duration of the pulse with similar time courses. Thus, three different ER-GEVIs derived from hVoS probes confirm previously reported responses of the ER membrane to plasma membrane voltage steps (16-18). The increase in fluorescence following a negative voltage step to the plasma membrane indicates that DPA is driven away from the mCeFP located on the cytoplasmic side of the ER membrane. Thus, a negative voltage step to the plasma membrane makes the cytoplasm more negative relative to the ER lumen, consistent with the sign of the voltage changes observed with other ER probes (17, 18).

The negative voltage steps employed in Figs. 2C and 2G will drive negatively charged DPA out of the cell, and if this reduces the cytosolic DPA concentration the fluorescence could increase. Although the flux of DPA should be very slow we felt it was important to test this possibility. This was done by first bathing cells in 8 μM DPA for 30 minutes and then transferring to a DPA-free bathing solution. Since cultured cells lose DPA at a rate of ∼ 1% per minute (38) this gives us an opportunity to record from a cell in the absence of extracellular DPA. We patch clamped cells within 10 minutes of transferring to the DPA-free solution and stepped the voltage from -70 to +80 mV. This would drive DPA into the cell if it were in the bathing solution but with a DPA-free solution a positive membrane potential cannot increase the intracellular DPA concentration. This positive voltage step elicited a clear reduction in fluorescence (Fig. 2H; this experiment was repeated in 4 cells). In the absence of extracellular DPA the fluorescence decrease cannot be attributed to an increase in cytosolic DPA concentration. We can thus conclude that it reflects DPA movement within the ER membrane driven by a change in Ψ_ER_.

### Spread of voltage change

We found that Ψ_ER_ does not change uniformly throughout the entire cell but spreads spatially over time. This is illustrated in an mCeFP-Sec61β-expressing cell in Fig. 3A with two succussive spatial maps of response magnitude, one early, 0.05-0.15 sec after the pulse start (Fig. 3B) and one later, 0.9-1.0 sec after the pulse start (Fig. 3C). The maps show that ER voltage in the lower pole of the cell (brown region in Fig. 3A) precedes the ER voltage change in a more central part of the cell (purple region in Fig. 3A). Normalized fluorescence traces from these two regions show a more rapid change at the pole (Fig. 3D, brown trace) and a slower change at the more central location (lavender trace). A second example from a different cell illustrates the same trend of nonuniform changes in Ψ_ER_, and with a more rapid change near a pole (Fig. 3E-3H).

**Figure 3.**
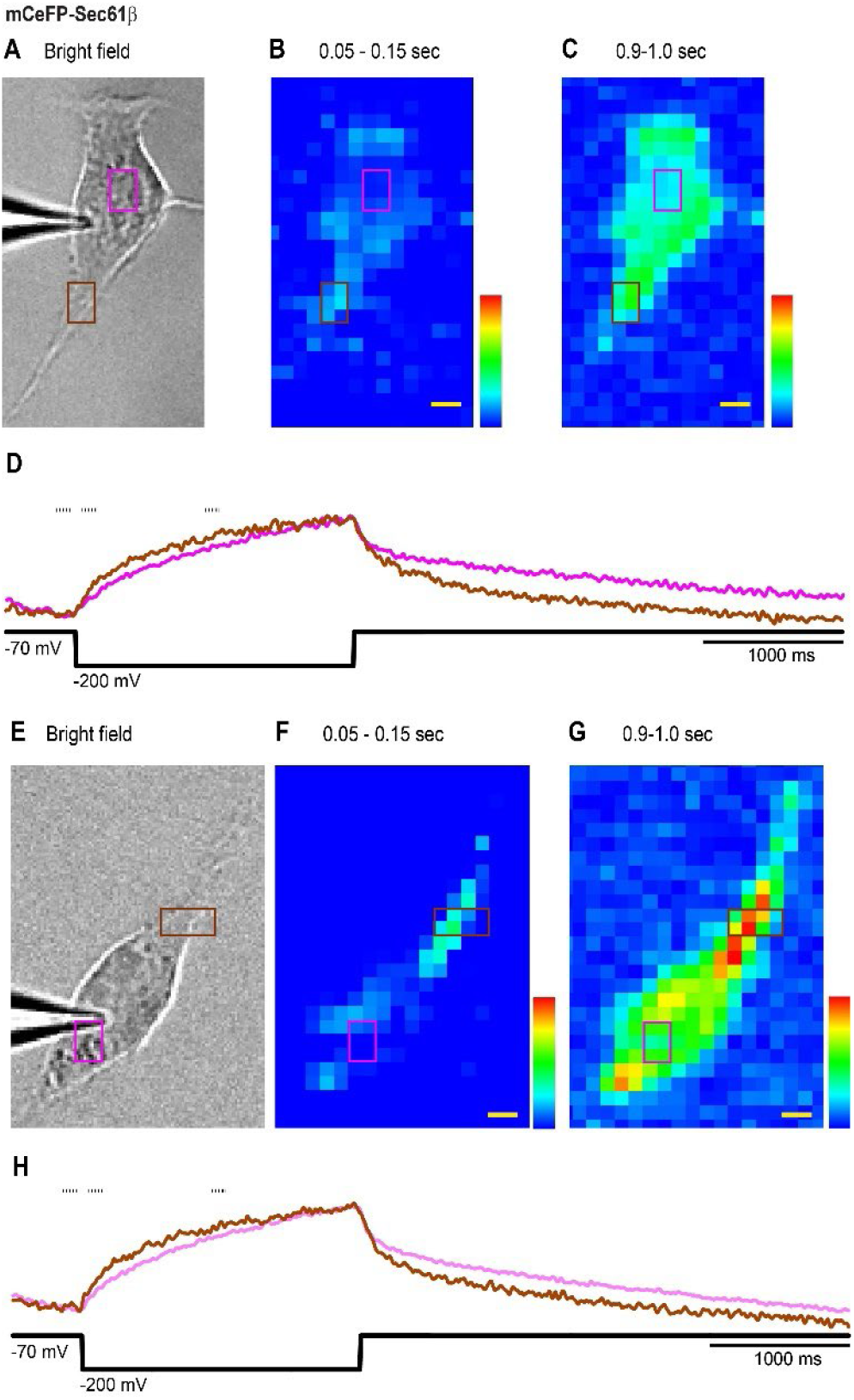
The dynamics of ER signals from different cellular locations. **A** and **E.** Bright field images of two patch-clamped HEK293T cells expressing mCeFP-Sec61β. **B** and **F**. Images early during a pulse from -70 to -200 mV. **C** and **G**. images later during the pulse. The times of the frames averaged to produce the images are indicated by the dashes above the fluorescence traces in D and H. The image before the start of the pulse was subtracted from images early during the voltage step (the first image, **B** and **F**) and later during the voltage step (the second image, **C** and **G**). **D** and **H**. The violet and brown traces represent the optical signals from the violet and brown rectangles indicated in (**A**-**C** and **E**-**G**). These traces have been normalized to their maxima in order to highlight their different time courses. Traces were low-pass filtered in Turbo-SM at a setting of 15. The black traces below indicate the voltage pulse protocols. Images were recorded at a frame rate of 200 Hz. The times above the images are with respect to the starts of the pulses. Scale bar = 5 μm.

### Voltage dependence

hVoS signals have a highly characteristic voltage dependence defined by the movement of DPA between the two faces of the membrane (19). We therefore applied a range of voltage steps as illustrated in Fig. 4A for CeFP-Sec61β. We measured the change from just before the start, to the very end of the pulse, and since the slow process is still incomplete this is a lower bound. We then plotted fluorescence versus voltage for all three ER-GEVIs as well as the plasma membrane probe hVoS3 for comparison (Fig. 4B). All the plots were well fitted by a Boltzmann function of the form

**Figure 4.**
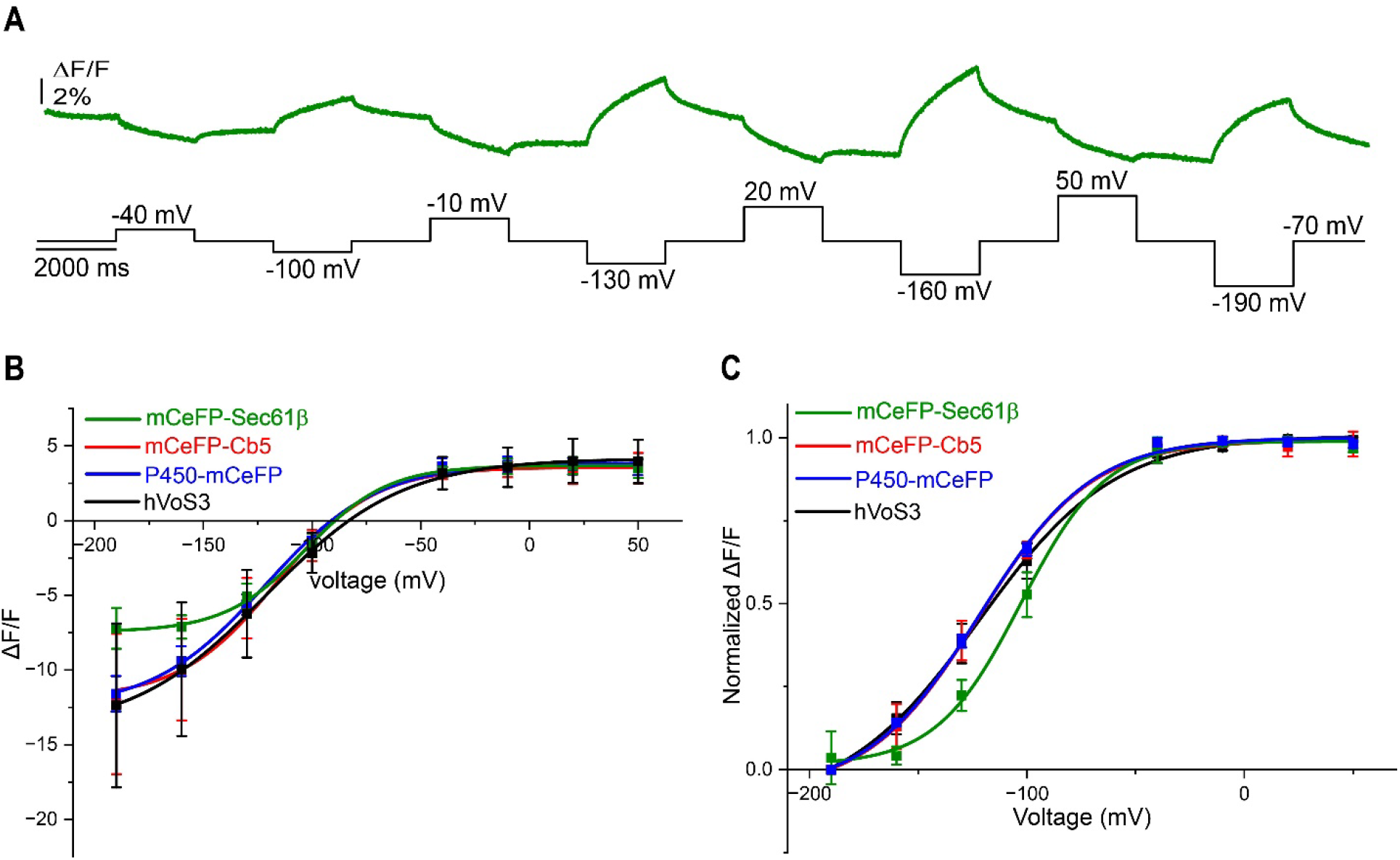
Voltage dependence of GEVI responses at the ER and plasma membrane. **A**. Trace of fluorescence from a mCeFP-Sec61β-expressing HEK293T cell with steps from -70 mV to different voltages (black trace below). **B**. Plots of fluorescence change (ΔF/F, from just before the start to the pulse end) versus voltage, together with Boltzmann fits (see text). **C**. Plots were normalized to their maximum and shifted to zero at *F*_*min*_ at the left extreme to compare the voltage dependence of the 4 probes. The green, red, blue and black curves are mCeFP-Sec61β, mCeFP-Cb5, P450-mCeFP and hVoS3, respectively. Data recorded at a frame rate of 200 Hz. N = 5 cells for each plot. Error bars are standard errors.

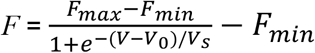

This function represents the voltage dependent distribution between two states of a charged molecule in a voltage gradient, and is widely used to characterize voltage gating of ion channels (39, 40). At the positive voltage limit with *F* = *F*_*max*_, all the DPA is on one side; at the negative limit with *F* = *F*_*min*_, all the DPA is on the other side. This equation thus expresses the voltage dependent distribution of DPA between the two faces of the membrane (19). *V*_*0*_ is the midpoint of the transition; *V*_*s*_ corresponds to the range over which the transition occurs and is referred to as the steepness parameter. The parameters from the fits are presented in Table 1. We normalized these plots in Fig. 4C and note that the normalized plots for mCeFP-Cb5 (red trace and points in Fig. 4C) and P450-mCeFP (blue) are nearly superimposable with the plot the plasma membrane probe hVoS3 (black). By contrast, the voltage dependence of mCeFP-Sec61β is right-shifted and is steeper (green). This may be an indication of subtle differences in the spatial distributions of the three constructs illustrated in Fig. 1, with P450-mCeFP and mCeFP-Cb5 targeting ER regions more strongly coupled to the plasma membrane. mCeFP-Sec61β then targets a region less tightly coupled, where the ER can escape the clamp at the plasma membrane. The greater steepness may reflect voltage-gated channels in this part of the ER that open as the voltage becomes more negative to take control of the voltage and move it to a level defined by the selectivity of the newly opened channels. Voltage-gated channels in the ER may also be responsible for the change in the slope of the fluorescence-voltage plot seen with ER-ASAP3 (18). Table 1 presents max-ΔF/F values (*F*_*max*_*-F*_*min*_) that represent the maximum fluorescence change that can be achieved between the low-voltage and high-voltage limits. These are important parameters for evaluating probe performance; their magnitudes reflect the limiting change in FRET between DPA and the fluorescent protein (31).

**Table 1.** Parameters for voltage dependence Parameters from fits of the Boltzmann function (see text) to the plots of Fig. 4B. Max ΔF/F was *F*_*max*_ - *F*_*min*_. Points are means ± standard errors. N is the number of experiments/cells.

| | $V_0$ (mV) | $V_s$ (mV) | Max-$\Delta F/F$ % | N |
| --- | --- | --- | --- | --- |
| mCeFP-Sec61β | $-102.7 \pm 1.5$ | $19.5 \pm 1.8$ | $11.1 \pm 0.27$ | 5 |
| mCeFP-Cb5 | $-122.0 \pm 2.4$ | $25.7 \pm 2.3$ | $15.5 \pm 1.20$ | 5 |
| P450-mCeFP | $-121.7 \pm 2.7$ | $26.0 \pm 2.6$ | $16.44 \pm 0.65$ | 5 |
| hVoS3 | $-121.9 \pm 2.3$ | $31.1 \pm 2.1$ | $18.23 \pm 0.49$ | 5 |

### Kinetics

For all three ER-GEVIs the fluorescence begins to change at the onset of a stepwise change in voltage, and gradually levels off over the course of the two-second pulse. To explore the dynamics of the apparent coupling between the plasma membrane and ER we fitted the fluorescence time course to exponential functions. For all three probes, a sum of two exponentials fitted the time courses very well, both for the on- and off-responses (Fig. 5).

**Figure 5.**
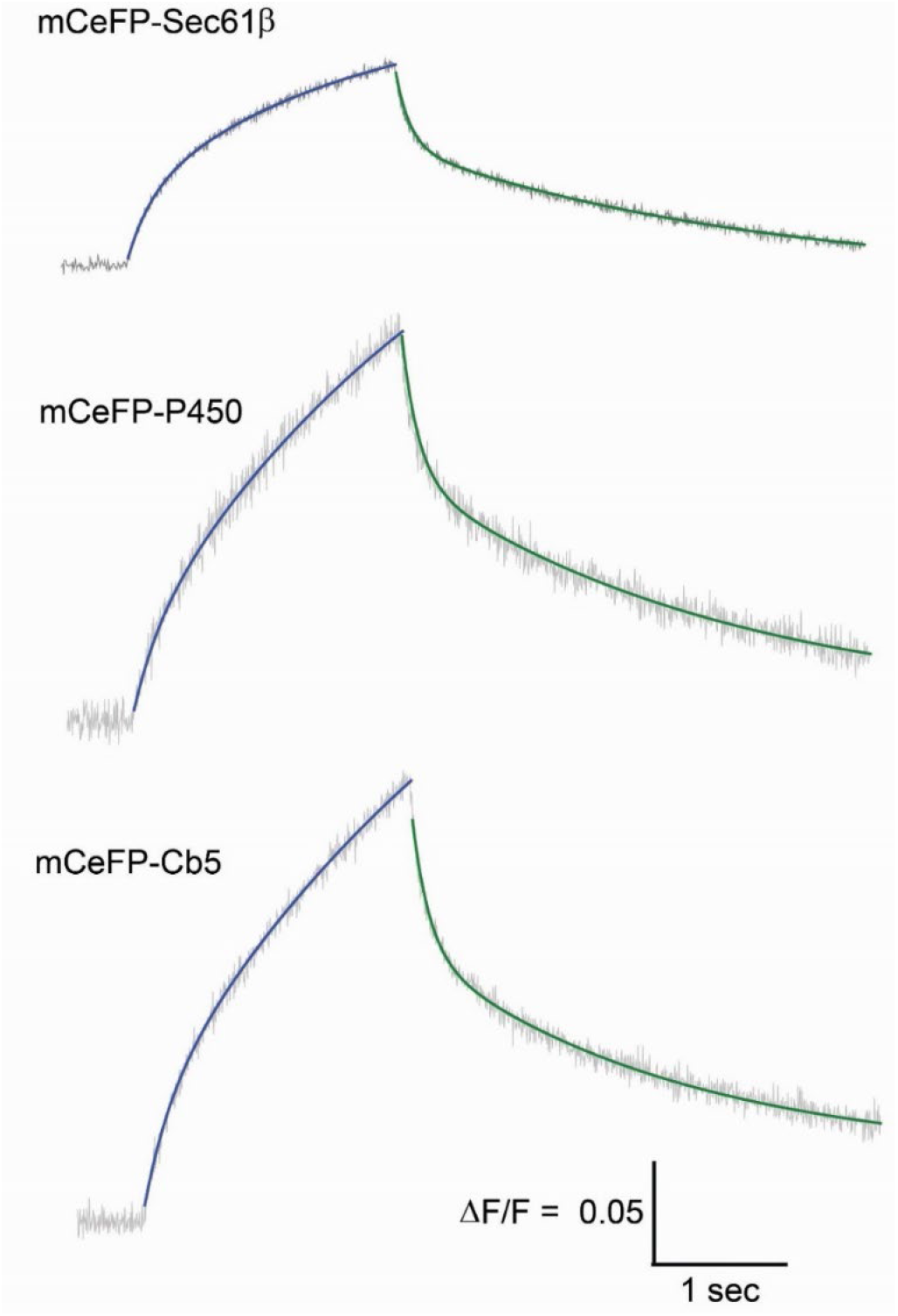
Double-exponential kinetics. Fits of the time course of mCeFP-Sec61β, mCeFP-P450, and mCeFP-Cb5 for voltage steps from -70 to -200 mV. A sum of two exponentials was fitted to both the on-responses (blue) andoff-responses (green); fluorescence is grey. The time constants and fraction of the fast component are presented in Fig. 6. The calibration bars apply to all three traces.

The time constants from the fits for all three ER-GEVIs are presented in Fig 6. Both on and off transitions have a fast component with a τ of ∼100-200 msec and a slow component with a τ of more than a second. ANOVA indicated that as a group the τ_on_-fast values are significantly longer than τ_off_-fast values and the fraction of the fast component (A_fast_/(A_fast_+A_slow_) is greater for off-responses. τ_fast_ and the fraction of fast component of the on-response of mCeFP-Sec61β are significantly different from the other probes, and there is a significant difference between the fraction of the fast component for the off-response between mCeFP-Sec61β and mCeFP-Cb5. The differences in charging kinetics may be yet another indication of spatial heterogeneity of expression between the compartments of ER targeted by the three probes.

**Fig. 6.**
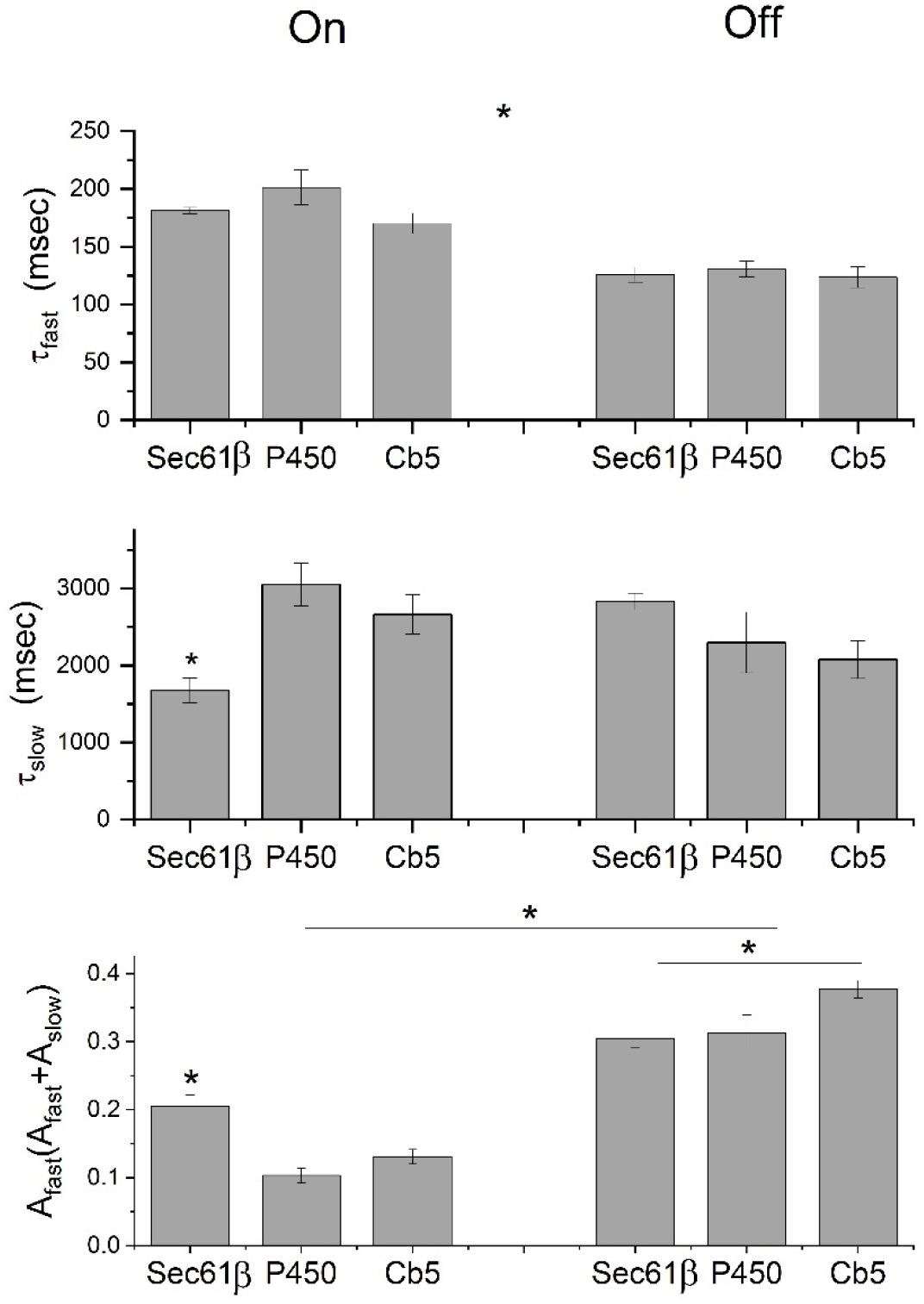
Time constants were determined from double exponential fits to the on-transition and the off-transition (from Fig. 5, green and blue, respectively). Means are presented ± standard error with N=5 experiments per cell for Sec61β and Cb5 and N=4 for P450. ^*^ denotes significant differences by ANOVA (P<0.05) for the indicated values or groups.

### Equivalent circuit

The dynamic changes in Ψ_ER_ reported by these three ER-GEVIs can be interpreted in terms of current passing through a resistance located at contacts between the plasma and ER membranes that connect the extracellular solution to the ER lumen. We therefore designed an equivalent circuit which captures some of the essential observations (Fig. 7). The left side of this circuit depicts the plasma membrane and voltage clamp, V_c_. This part corresponds to the generally accepted circuit used to interpret whole-cell patch-clamp recording (37). A voltage step charges the plasma membrane with a time constant of R_S_C_PM_ (provided that R_PM_ >> R_S_), where R_S_ is the series or access resistance of the patch electrode and C_PM_ is the capacitance of the plasma membrane, determined by transient cancelation (37). R_S_C_PM_ is typically ∼100 μsec. Because R_PM_ >> R_S_ the voltage across the plasma membrane is nearly equal to V_c_. Because the plasma membrane charges much more rapidly than the ER membrane, we can see by analogy that a similar dynamic situation applies to the charging of the ER. Charging the plasma membrane drives current through a resistor R_PM-ER_ which we hypothesize couples the plasma and ER membranes. This current charges the ER membrane capacitance, C_ER_. Because the plasma membrane charges so much more rapidly than the ER membrane, we can approximate the time constant for ER charging as R_PM-ER_C_ER_. This is analogous to the R_S_C_PM_ time constant for plasma membrane charging. As noted above, the expression for the plasma membrane charging time constant of R_S_C_PM_ depends on the condition R_PM_>>R_S_. Likewise, the expression for the ER membrane charging time constant of R_PM-ER_C_ER_ depends on the condition that R_ER_>>R_PM-ER_. The similarity in plots of fluorescence versus voltage for hVoS3 and two of the ER-GEVIs (Fig. 4) supports this condition.

**Figure 7.**
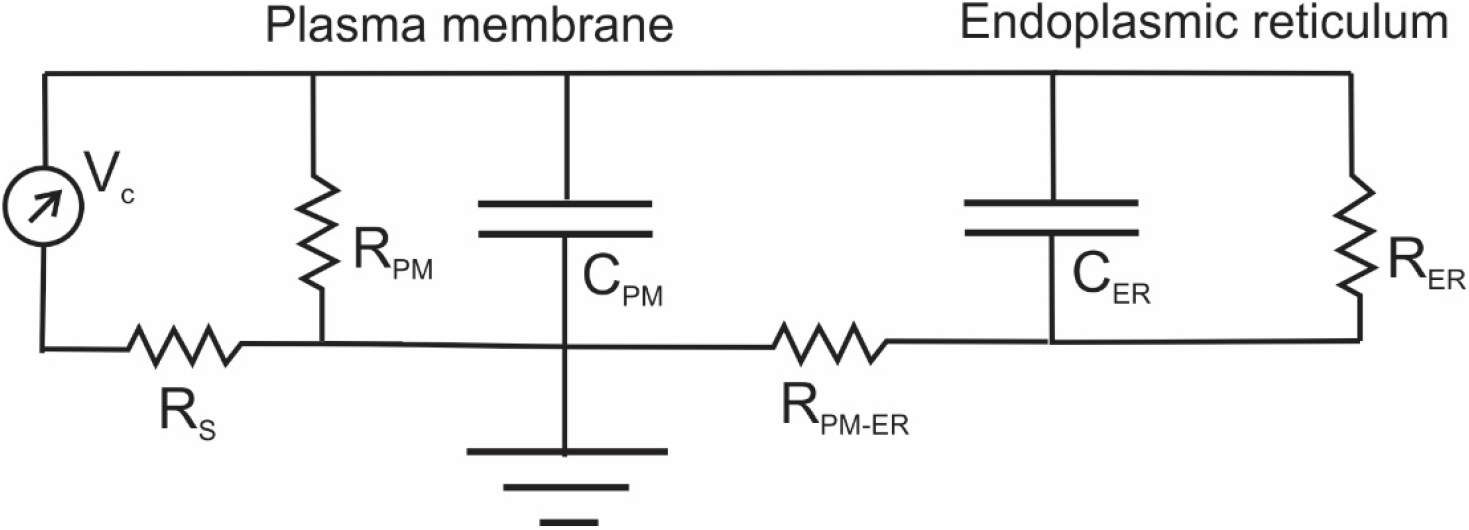
Equivalent circuit. The plasma membrane, ER membrane, and their electrical coupling are represented by an equivalent circuit. The voltage clamp (V_c_) on the left applies a voltage step to the plasma membrane. The plasma membrane has a resistance R_PM_ and capacitance C_PM_; the series resistance of the electrode is R_S_; C_PM_ is charged with a time constant of R_S_C_PM_, typically ∼100 μsec (37). The ER has a resistance R_ER_ and a capacitance C_ER_, and the plasma membrane-ER contact presents a resistance R_PM-ER_ to current flow to the ER lumen. Because the charging of C_ER_ is much slower than the charging of C_PM_, the ER charging can be treated as a dynamically separate process with a time constant of R_PM-ER_C_ER_.

We can estimate C_ER_ from the knowledge that the ER membrane within an epithelial cell has an area 25 times larger than the area of the plasma membrane (41). We measured the capacitance of HEK293T cells by transient cancelation (37) as 13.0 ± 2.2 pF (mean ± standard deviation, N=11), so C_ER_ is 25 times larger or 325 pF. This value and the charging time constant can be used to calculate R_PM-ER_. The ER charges in two phases (Fig. 5), and we chose to focus on the faster component because the slower component may reflect equalization within a complex geometry (37, 42, 43). The τ_on_-fast values in Table 2 are similar and their average is 180 msec. Using this and the above value for C_ER_ we obtain R_PM-ER_ = 550 MΩ (conductance = 1.82 nS). If the contact between the plasma membrane and ER contains gap junction-like channels spanning two lipid bilayers, then we can estimate the number of channels from our value of R_PM-ER_. The single channel conductances of connexins range from 22 – 310 pS but most are around 100 pS (44). Our estimate of R_PM-ER_ is compatible with 18 channels with a conductance of 100 pS. One scenario is 18 separate ER-plasma membrane contact sites within a cell, each with a single gap junction like channel. A channel found in ER contacts is the calcium release activated channel (CRAC), for which a conductance of 0.7 pS has been determined (45). If a channel with this conductance mediated ER-plasma membrane coupling, then over 2000 of them would be needed for our estimate of R_PM_-_ER_. We note that this estimate assumes that ER charging is passive, without an influence from the gating of channels in the ER membrane. This is consistent with the voltage dependence of mCeFP-Cb5 and P450-mCeFP, but as noted above the voltage dependence of mCeFP-Sec61β suggests that it targets ER that is influenced by voltage-gated channels (Fig. 3C and Table 1).

## Discussion

The ER-GEVIs developed in this study generate fluorescence signals that track changes in Ψ_ER_. The probes exhibit a clear cytosolic localization, colocalize with an established ER marker, and employ well-established ER targeting motifs from Sec61β (34), P450 (27), and Cb5 (28-30). All three probes show no significant labeling of the plasma membrane. Their fluorescence changes following voltage steps to the plasma membrane are much slower than the changes seen with a plasma membrane GEVI, confirming their exclusion from the cell surface. The fluorescence changes depended on the presence of DPA (Fig. 2C), indicating that these probes sense Ψ_ER_ and not another property such as pH or metabolite level. Furthermore, a positive voltage step applied to a DPA-loaded cell produced a fluorescence decrease in a DPA-free bathing solution (Fig. 2H), indicating that the signal does not reflect a voltage-induced change in cytosolic DPA concentration. Thus, these hVoS-based probes act as ER-GEVIs and sense the voltage at the ER membrane, Ψ_ER_.

Changing the voltage across the plasma membrane in the negative direction, to make the cytoplasm more negative relative to the extracellular space, increased the fluorescence emission of our probes. All three probes place their fluorescent protein on the cytoplasmic face of the ER membrane, so the increase in fluorescence means that negatively charged DPA moves within the ER membrane, away from the cytoplasmic face and toward the luminal face. This indicates that making the cytoplasm more negative with respect to the extracellular space also makes the cytoplasm more negative with respect to the ER lumen. This confirms reports with other ER voltage sensors regarding the relation between the plasma membrane potential and Ψ_ER_ (17, 18). This relation, along with the dynamics of the signals, indicates that there is electrical continuity between the ER lumen and extracellular space. A voltage change at the plasma membrane is not transmitted to the ER membrane indirectly through ionic or chemical changes in the cytoplasm. A buildup of cytosolic Ca^2+^ or protons, or the generation of a metabolite by a voltage dependent enzyme at the plasma membrane could not initiate a change Ψ_ER_ this rapidly.

Experiments in other labs with different ER-GEVIs have shown that voltage steps to the plasma membrane produce rapid changes in Ψ_ER_ (16-18), but the kinetics of these responses were not quantified. In one of these studies (16) the traces display three kinetic components, one rapid component from the plasma membrane and two slower components qualitatively similar to those reported here. In the other studies (17, 18) the display scales did not render the time course and we cannot evaluate response kinetics. We note that charging on the order of >100 msec cannot reflect heterogeneous charging of the plasma membrane. The assumption of isopotentiality is embedded in the equivalent circuit of the whole-cell patch-clamp which quantitatively accounts for the rapid charging transients generally observed in geometrically simple cells (37). Local voltage equalization is ∼ 1,000 time faster than the cell membrane time constant, which is on the order of milliseconds (46). In geometrically complex neurons with processes the charging transients can be as long as 10 msec (37, 42, 43). Thus, the time constants for Ψ_ER_ charging that range from 170 to 3000 msec are far too long to reflect plasma membrane charging.

It is notable that probes incorporating three different ER targeting motifs produce qualitatively similar signals. Given the similar slow dynamics with three different ER-GEVIs, it is difficult to see how the targeting motif could influence response kinetics. The fluorescence changes depend on the presence of DPA (Fig. 2C) and the voltage dependent movement of DPA is known to be fast (38). If voltage acted on the targeting motif to perturb the fluorescent protein, we would not expect to this perturbation to be similar with three different targeting motifs, and the effect would not depend on DPA. If DPA interacts with the targeting motif in a voltage dependent manner this interaction would have to overwhelm the FRET interaction with the free DPA in the lipid bilayer. However, there were quantitative differences in probe voltage dependence (Table 1), with a greater steepness for mCeFP-Sec61β (Fig. 4C). There were also quantitative differences in their charging kinetics (Fig. 6). Furthermore, although the probes showed high colocalization with mCherry-Sec61β, there were small but significant differences (Fig. 1I). These results point to a non-uniform distribution of these probes within the ER. The Sec61 complex functions in protein synthesis and is concentrated in the rough ER, whereas the other probes should be distributed through both the rough and smooth ER. Structural studies have revealed that the ER has distinct subdomains (1, 41, 47). The spatial dynamics of charging suggest that ER domains closer to the cell periphery may charge more rapidly than deeper compartments (Fig. 3). The extent of the voltage change can vary as well as the speed, but this will be difficult to assess due to the difficulty of resolving variations in probe expression levels and background fluorescence arising from both cell autofluorescence and mistargeted probe. The different response properties of these three probes thus support the view that the ER is heterogeneous in its voltage signaling, as suggested by Campbell et al (18). The ER membrane harbors a number of ion channels (13), and the steeper voltage dependence of mCeFP-Sec61β responses could reflect the presence of voltage-gated channels in the ER compartment targeted by this probe. The voltage change would then open channels which would take control of the voltage across the membrane compartment in which they reside. This supports the suggestion that ion channels can control the spread of voltage within the ER (18).

The responses of these ER-GEVIs started at the onset of the voltage step applied to the plasma membrane and rose slowly toward a plateau. Thus, a voltage step to the plasma membrane initiates a time-dependent charging of the ER membrane. We interpreted this in terms of an equivalent circuit with a charging time constant of R_PM-ER_C_ER_ (Fig. 7). R_PM-ER_ could reflect conduction through channels at plasma membrane-ER contact sites that span the two membranes. Plasma membrane-ER contacts have received a great deal of attention (48), and these sites mediate the replenishment of ER Ca^2+^ stores (49-51). Ion conduction through the STIM1-ORAI1 complex may account for all or part of R_PM-ER_ depicted in Fig. 7. ER Ca^2+^ stores can be replenished without a detectible change in cytosolic Ca^2+^ concentration (52), and a channel connection through both membranes from the extracellular medium to the ER lumen could accomplish this.

The ER harbors a host of ion channels that have the capacity to generate voltage and mediate ion fluxes (13). ER Ca^2+^ signaling (2, 7) will be modulated by Ψ_ER_. The ability to track Ψ_ER_ in real time with sub-cellular spatial resolution will enable the use of these ER-GEVIs to study vital ER functions at a new level of detail. Such applications may be hampered by spatial variations in probe expression and DPA distribution, and selecting an appropriate ER-GEVI may depend on the function to be studied. Spillover of probes into other cytoplasmic compartments may complicate such studies, and future efforts to assess these pitfalls will be important in the effective use of ER-GEVIs.

ER-GEVIs such as those introduced here provide an experimental window into dynamic changes in Ψ_ER_. Optical monitoring of Ψ_ER_ promises to advance the study of ER processes such as Ca^2+^ signaling, lipid biosynthesis, and protein synthesis. We have shown that these probes can be used to assess the electrical coupling between the ER and plasma membranes. This approach will enable the identification and characterization of the proteins that mediate and control this coupling. Experiments with these probes in neurons with complex morphology have the potential to reveal specialized roles for Ψ_ER_ signaling in neuronal somata, dendrites, and synaptic terminals. ER dysfunction is implicated in many diseases including neurodegeneration and lipid storage disorders, and tracking Ψ_ER_ could provide valuable insights into these pathological conditions. Finally, using ER-GEVIs for drug screening could help to identify therapeutic compounds that modulate Ψ_ER_ and ER-plasma membrane coupling.

## Author contributions

MSR performed all the experiments and generated all the reagents. Both authors designed experiments, analyzed data, and wrote the manuscript.

The Wisconsin Alumni Research Foundation has patented these probes – US 11,846,628 B2.

## Acknowledgements

Supported by NIH grant R35 NS127219

## References

1. Baumann, O., and B. Walz. 2001. Endoplasmic reticulum of animal cells and its organization into structural and functional domains. Int Rev Cytol 205:149–214.

2. Schwarz, D. S., and M. D. Blower. 2016. The endoplasmic reticulum: structure, function and response to cellular signaling. Cell Mol Life Sci 73:79–94.

3. Boelens, J., S. Lust, F. Offner, M. E. Bracke, and B. W. Vanhoecke. 2007. Review. The endoplasmic reticulum: a target for new anticancer drugs. In Vivo 21:215–226.

4. Stutzmann, G. E., and M. P. Mattson. 2011. Endoplasmic reticulum Ca(2+) handling in excitable cells in health and disease. Pharmacol Rev 63:700–727.

5. Gallo, A., C. Vannier, and T. Galli. 2016. Endoplasmic Reticulum-Plasma Membrane Associations:Structures and Functions. Annu Rev Cell Dev Biol 32:279–301.

6. Valm, A. M., S. Cohen, W. R. Legant, J. Melunis, U. Hershberg, E. Wait, A. R. Cohen, M. W. Davidson, E. Betzig, and J. Lippincott-Schwartz. 2017. Applying systems-level spectral imaging and analysis to reveal the organelle interactome. Nature 546:162–167.

7. Berridge, M. J. 2002. The endoplasmic reticulum: a multifunctional signaling organelle. Cell Calcium 32:235–249.

8. Raymond, C. R. 2007. LTP forms 1, 2 and 3: different mechanisms for the “long” in long-term potentiation. Trends Neurosci 30:167–175.

9. Padamsey, Z., W. J. Foster, and N. J. Emptage. 2019. Intracellular Ca(2+) Release and Synaptic Plasticity: A Tale of Many Stores. Neuroscientist 25:208–226.

10. Bezprozvanny, I., and E. T. Kavalali. 2020. Presynaptic endoplasmic reticulum and neurotransmission. Cell Calcium 85:102133.

11. Berridge, M. J. 2007. Inositol trisphosphate and calcium oscillations. Biochem Soc Symp:1-7.

12. Lewis, R. S. 2003. Calcium oscillations in T-cells: mechanisms and consequences for gene expression. Biochem Soc Trans 31:925–929.

13. Takeshima, H., E. Venturi, and R. Sitsapesan. 2015. New and notable ion-channels in the sarcoplasmic/endoplasmic reticulum: do they support the process of intracellular Ca(2)(+) release? J Physiol 593:3241–3251.

14. Castillo-Velasquez, C., E. Matamala, D. Becerra, P. Orio, and S. E. Brauchi. 2024. Optical recordings of organellar membrane potentials and the components of membrane conductance in lysosomes. J Physiol 602:1637–1654.

15. Matamala, E., C. Castillo, J. P. Vivar, P. A. Rojas, and S. E. Brauchi. 2021. Imaging the electrical activity of organelles in living cells. Commun Biol 4:389.

16. Sepehri Rad, M., L. B. Cohen, O. Braubach, and B. J. Baker. 2018. Monitoring voltage fluctuations of intracellular membranes. Sci Rep 8:6911.

17. Klier, P. E. Z., R. Roo, and E. W. Miller. 2022. Fluorescent indicators for imaging membrane potential of organelles. Curr Opin Chem Biol 71:102203.

18. Campbell, E. P., A. A. Abushawish, L. A. Valdez, M. K. Bell, M. Haryono, P. Rangamani, and B. L. Bloodgood. 2023. Electrical signals in the ER are cell type and stimulus specific with extreme spatial compartmentalization in neurons. Cell Rep 42:111943.

19. Chanda, B., R. Blunck, L. C. Faria, F. E. Schweizer, I. Mody, and F. Bezanilla. 2005. A hybrid approach to measuring electrical activity in genetically specified neurons. Nat Neurosci 8:1619–1626.

20. Bayguinov, P. O., Y. Ma, Y. Gao, X. Zhao, and M. B. Jackson. 2017. Imaging Voltage in Genetically Defined Neuronal Subpopulations with a Cre Recombinase-Targeted Hybrid Voltage Sensor. J Neurosci 37:9305–9319.

21. Ma, Y., P. O. Bayguinov, and M. B. Jackson. 2017. Action potential dynamics in fine axons probed with an axonally-targeted optical voltage sensor. eNeuro (in press).

22. Scheuer, K. S., A. M. Jansson, M. Shen, X. Zhao, and M. B. Jackson. 2025. Fxr1 Deletion from Cortical Parvalbumin Interneurons Modifies Their Excitatory Synaptic Responses. eNeuro 12.

23. Chertkova, A. O., M. Mastop, M. Postma, N. van Bommel, S. van der Niet, K. L. Batenburg, L. Joosen, T. W. J. Gadella, Y. Okada, and J. Goedhart. 2020. Robust and Bright Genetically Encoded Fluorescent Markers for Highlighting Structures and Compartments in Mammalian Cells. bioRxiv:160374.

24. Linxweiler, M., B. Schick, and R. Zimmermann. 2017. Let’s talk about Secs: Sec61, Sec62 and Sec63 in signal transduction, oncology and personalized medicine. Signal Transduct Target Ther 2:17002.

25. Greenfield, J. J., and S. High. 1999. The Sec61 complex is located in both the ER and the ER-Golgi intermediate compartment. J Cell Sci 112 (Pt 10):1477–1486.

26. Matsuura, S., Y. Fujii-Kuriyama, and Y. Tashiro. 1978. Immunoelectron microscope localization of cytochrome P-450 on microsomes and other membrane structures of rat hepatocytes. J Cell Biol 78:503–519.

27. Szczesna-Skorupa, E., and B. Kemper. 1993. An N-terminal glycosylation signal on cytochrome P450 is restricted to the endoplasmic reticulum in a luminal orientation. J Biol Chem 268:1757–1762.

28. D’Arrigo, A., E. Manera, R. Longhi, and N. Borgese. 1993. The specific subcellular localization of two isoforms of cytochrome b5 suggests novel targeting pathways. J Biol Chem 268:2802–2808.

29. Mitoma, J., and A. Ito. 1992. The carboxy-terminal 10 amino acid residues of cytochrome b5 are necessary for its targeting to the endoplasmic reticulum. EMBO J 11:4197–4203.

30. Zhu, W., A. Cowie, G. W. Wasfy, L. Z. Penn, B. Leber, and D. W. Andrews. 1996. Bcl-2 mutants with restricted subcellular location reveal spatially distinct pathways for apoptosis in different cell types. EMBO J 15:4130–4141.

31. Wang, D., Z. Zhang, B. Chanda, and M. B. Jackson. 2010. Improved probes for hybrid voltage sensor imaging. Biophys J 99:2355–2365.

32. Markwardt, M. L., G. J. Kremers, C. A. Kraft, K. Ray, P. J. Cranfill, K. A. Wilson, R. N. Day, R. M. Wachter, M. W. Davidson, and M. A. Rizzo. 2011. An improved cerulean fluorescent protein with enhanced brightness and reduced reversible photoswitching. PLoS One 6:e17896.

33. Lee, S., T. Geiller, A. Jung, R. Nakajima, Y. K. Song, and B. J. Baker. 2017. Improving a genetically encoded voltage indicator by modifying the cytoplasmic charge composition. Sci Rep 7:8286.

34. Zurek, N., L. Sparks, and G. Voeltz. 2011. Reticulon short hairpin transmembrane domains are used to shape ER tubules. Traffic 12:28–41.

35. Mehta, S., N. N. Aye-Han, A. Ganesan, L. Oldach, K. Gorshkov, and J. Zhang. 2014. Calmodulin-controlled spatial decoding of oscillatory Ca2+ signals by calcineurin. Elife 3:e03765.

36. Cho, K. F., T. C. Branon, S. Rajeev, T. Svinkina, N. D. Udeshi, T. Thoudam, C. Kwak, H. W. Rhee, I. K. Lee, S. A. Carr, and A. Y. Ting. 2020. Split-TurboID enables contact-dependent proximity labeling in cells. Proc Natl Acad Sci U S A 117:12143–12154.

37. Marty, A., and E. Neher. 1995. Tight-seal whole-cell recording. In Single-Channel Recording. B. Sakmann, and E. Neher, editors. Plenum, New York. 31–51.

38. Oberhauser, A. F., and J. M. Fernandez. 1995. Hydrophobic ions amplify the capacitive currents used to measure exocytotic fusion. Biophys J 69:451–459.

39. Bezanilla, F. 2000. The voltage sensor in voltage-dependent ion channels. Physiol Rev 80:555–592.

40. Jackson, M. B. 2006. Molecular and cellular biophysics. Cambridge University Press, Cambridge ; New York.

41. Weibel, E. R., W. Staubli, H. R. Gnagi, and F. A. Hess. 1969. Correlated morphometric and biochemical studies on the liver cell. I. Morphometric model, stereologic methods, and normal morphometric data for rat liver. J Cell Biol 42:68–91.

42. Jackson, M. B. 1992. Cable analysis with the whole-cell patch clamp. Theory and experiment. Biophys J 61:756–766.

43. Rall, W. 1969. Distributions of potential in cylindrical coordinates and time constants for a membrane cylinder. Biophys J 9:1509–1541.

44. Bennett, M. V., and V. K. Verselis. 1992. Biophysics of gap junctions. Semin Cell Biol 3:29–47.

45. Prakriya, M., and R. S. Lewis. 2006. Regulation of CRAC channel activity by recruitment of silent channels to a high open-probability gating mode. J Gen Physiol 128:373–386.

46. Rall, W. 1969. Time constants and electrotonic length of membrane cylinders and neurons. Biophys J 9:1483–1508.

47. Borgese, N., M. Francolini, and E. Snapp. 2006. Endoplasmic reticulum architecture: structures in flux. Curr Opin Cell Biol 18:358–364.

48. Stefan, C. J., A. G. Manford, and S. D. Emr. 2013. ER-PM connections: sites of information transfer and inter-organelle communication. Curr Opin Cell Biol 25:434–442.

49. Liou, J., M. L. Kim, W. D. Heo, J. T. Jones, J. W. Myers, J. E. Ferrell, Jr., and T. Meyer. 2005. STIM is a Ca2+ sensor essential for Ca2+-store-depletion-triggered Ca2+ influx. Curr Biol 15:1235–1241.

50. Prakriya, M., S. Feske, Y. Gwack, S. Srikanth, A. Rao, and P. G. Hogan. 2006. Orai1 is an essential pore subunit of the CRAC channel. Nature 443:230–233.

51. Zhang, S. L., Y. Yu, J. Roos, J. A. Kozak, T. J. Deerinck, M. H. Ellisman, K. A. Stauderman, and M. D. Cahalan. 2005. STIM1 is a Ca2+ sensor that activates CRAC channels and migrates from the Ca2+ store to the plasma membrane. Nature 437:902–905.

52. Petersen, O. H., R. Courjaret, and K. Machaca. 2017. Ca(2+) tunnelling through the ER lumen as a mechanism for delivering Ca(2+) entering via store-operated Ca(2+) channels to specific target sites. J Physiol 595:2999–3014.

